# Loss of the Heparan Sulfate Proteoglycan Glypican5 facilitates long range Shh signaling

**DOI:** 10.1101/448621

**Authors:** Wei Guo, Henk Roelink

## Abstract

As a morphogen, Sonic Hedgehog (Shh) mediates signaling at a distance from its sites of synthesis. After secretion, Shh must traverse a distance through the extracellular matrix (ECM) to reach the target cells and activate the Hh response. Extracellular matrix proteins, in particular the Heparan Sulfate Proteoglycans (HSPGs) of the Glypican family have both negative and positive effects on non-cell autonomous Shh signaling, all attributed to their ability to bind Shh. Using mouse embryonic stem cell-derived mosaic tissues with compartments that lack the glycosyltransferases Exostosin1 (Ext1) and Exostosin2 (Ext2), or the HSPG core protein *Glypican5* we show that cells surrounded by a mutated extracellular matrix are highly proficient distributing Shh. In contrast, cells that lack *Ext1* function poorly secrete Shh. Our results confirm earlier observations that HSPGs can have both positive (Shh export) and negative influences (Shh distribution), and are supporting a model in which Shh presented on the cell surface in the context of HSPGs preferentially distributes into ECM that lacks HSPGs, possibly due to the absence of Shh sequestering molecules.

---

Sonic Hedgehog (Shh) and its paralog Desert Hh (Dhh) and Indian Hh Ihh can function as morphogens, signaling molecules that are produced locally and form a concentration gradient as spreading through surrounding tissue. The graded signal is interpreted by cells, in a dosage-dependent manner, to control gene expression and cell fate specification. Hhs are essential for patterning and differentiation in most animals (Briscoe and Thérond, 2013) and aberrant regulation of Hh pathway is associated with congenital anomalies, such as holoprosencephaly, and cancer (Hui and Angers, 2011; Jiang and Hui, 2008). The importance of a graded Hh concentration in tissue patterning during embryogenesis has long been recognized. It is, however, poorly understood how these gradients are established in the extracellular space.

Hh is synthesized as a precursor protein and undergoes autoproteolysis during which C-terminal domain is autocatalytically cleaved (Porter et al., 1995) and the remaining N-terminal domain (HhNp) is covalently modified by cholesterol at the C terminus and palmitoylated at the N-terminus (Bumcrot et al., 1995; Pepinsky et al., 1998). The addition of cholesterol and palmitic acid promotes the association of HhNp with sterol-rich membrane microdomains (Chen et al., 2004; Rietveld et al., 1999). Subsequent release of HhNp into the extracellular space requires the RND antiporter Dispatched1 (Disp1) (Burke et al., 1999), the CUB domain protein Scube2 (Creanga et al., 2012; Jakobs et al., 2014), and members of the ADAM family of sheddases (Damhofer et al., 2015; Dierker et al., 2009).

In the extracellular space Shh encounters extracellular matrix components that shape Hh gradients including heparan sulfate proteoglycans (HSPGs). HSPGs consist of a protein core (such as glypican and syndecan) to which heparan sulfate glycosaminoglycan chains are attached (Merton Bernfield et al., 1999). Heparan sulfate glycosaminoglycan chains are added to a core protein by the sequential action of individual glycosyltransferases and modification enzymes, in a three-step process involving chain initiation, polymerization, and modification (Yan and Lin, 2009). Ext1 and Ext2 form heteromeric complex of the glycosyltransferases and catalyze HS chain polymerization (Senay et al., 2000). Previous study showed that a gene trap mutation of *Ext1* resulted in substantial reduction of HS chain length (Österholm et al., 2009).

Genetic screens in *Drosophila* have shown that mutation of Ext1/2 orthologs (tout-velu, and brother of tout-velu), or the proteoglycans core proteins, the glypicans dally and dally-like protein impede Hh spread and reduce signaling range (Bellaiche et al., 1998; Han et al., 2004; Lin, 2004), indicating role for modified Glypicans facilitating Hh distribution. However, reduced HS synthesis in mice carrying a hypomorphic mutation in *Ext1*, results in an elevated range of Ihh signaling during embryonic chondrocyte differentiation (Koziel et al., 2004) which would be consistent with a negative activity of HSPG on Hh distribution. Glypicans can act either to sequester Hh ligand thus inhibiting signaling signaling or, in other cases, glypicans can stabilize the association of the ligand with the Ptc receptor to promote signaling (Filmus and Capurro, 2014; Ramsbottom and Pownall, 2016). For instance, Glypican3 (Gpc3) acts as a negative regulator of Shh activity by competing with Ptch1 for Shh binding (Capurro et al., 2008), whereas Gpc5 has been identified as a Shh co-receptor and promote downstream Shh signaling (Li et al., 2011; Witt et al., 2013).

As a long-range morphogen, Shh directs the pattern of neurogenesis by conferring positional information to ventral neural progenitors through a well-studied transcriptional response (Dessaud et al., 2008). Shh is produced ventrally from the notochord and floor plate, yielding a concentration gradient along the dorsoventral (DV) axis of the neural tube where ventral cell types have a high level of Shh pathway activation. These signaling events can be modeled by in vitro differentiating mESCs into neuralized embryoid bodies (nEBs) under neural induction conditions, which will be useful for studying the mechanism of morphogen action in a 3D environment (Meinhardt et al., 2014; Roberts et al., 2016; Wichterle et al., 2002). In the present study, in order to explore the role of Ext1/2 and glypicans in Shh distribution and pathway regulation, mosaic nEBs are generated consisting of various genome edited mESC lines as receiving cells and low number of Shh-expressing cells as Shh source. We show that absence of *Ext1/2* or *Glypican5* in surrounding cells increases the amount of extracellular Shh and enhances long-range Shh signaling. Our results demonstrate that HS-modified Gpc5 is a general inhibitor of Shh distribution, and its loss leads to activated Hh response. These results provide a simple explanation for the tumor suppressing activities of Ext1/2 and Glypican5, as their loss facilitates Shh signaling.

## Results

### Loss of *Ext1/2* in surrounding cells results in Strong Shh accumulation in nEBs

The use of mosaic nEBs has allowed us to precisely delineate roles of cells in the production of, transport of, and the response to Shh (Roberts et al., 2016). To assess if HSPGs regulate Shh distribution, we made genome edited mESC lines carrying homozygous null mutations in *Ext1 and Ext2*. The *Ext1* null were identified by analysis of the genomic loci, and further confirmed by the absence of *Ext1* transcripts (Sup). Mosaic nEBs comprised of majority of *Ext1*^−/−^ or *Ext2*^−/−^ cells and 1% wild type cells harboring the *EF1α:Shh* transgene (Roberts et al., 2016) provides us an in vitro model in which we can assess the non-cell autonomous effects of *Ext1/2* and measure the effect on the Hh response in recipient cells. The inclusion of small number of Shh expressing cells serves as sparse and localized sources of Shh in mosaic nEBs.

We first assayed the influence of *Ext1/2* nulls on Shh distribution by generating mosaic nEBs comprised of a large majority of *Shh*^−/−^; *Ptch1*^+/LacZ^; *Ext1*^−/−^ and *Shh*^−/−^; *Ptch1*^+/LacZ^; *Ext2*^−/−^ cells as responding cells and 3% Shh-expressing (otherwise wild type) cells as Shh source. Live staining with the anti-Shh monoclonal antibody 5E1 is expected to exclusively bind Shh present in the extracellular space, and Shh was not detected in nEBs without Shh expressing cells regardless of genotype (Fig. 1 A-C). We found small patches of extracellular Shh around the small number of Shh source cells present in the mosaic *Shh*^−/−^; *Ptch1*^+/LacZ^ nEBs (Fig. 1D). Surprisingly, the domains we detected Shh in *Ext1* and *Ext2* null nEBs were much larger than those in nEBs where the bulk of the cells is *Shh*^−/−^; *Ptch1*^+/LacZ^ (Fig. 1 E, F, quantified in Fig. 1G). This demonstrates that HSPGs negatively affect Shh distribution away from the sites of synthesis. It also indicates that Shh synthesized in cells with a HSPG competent matrix, nevertheless distributes preferentially into the matrix that lacks HSPGs.

**Figure 1.**
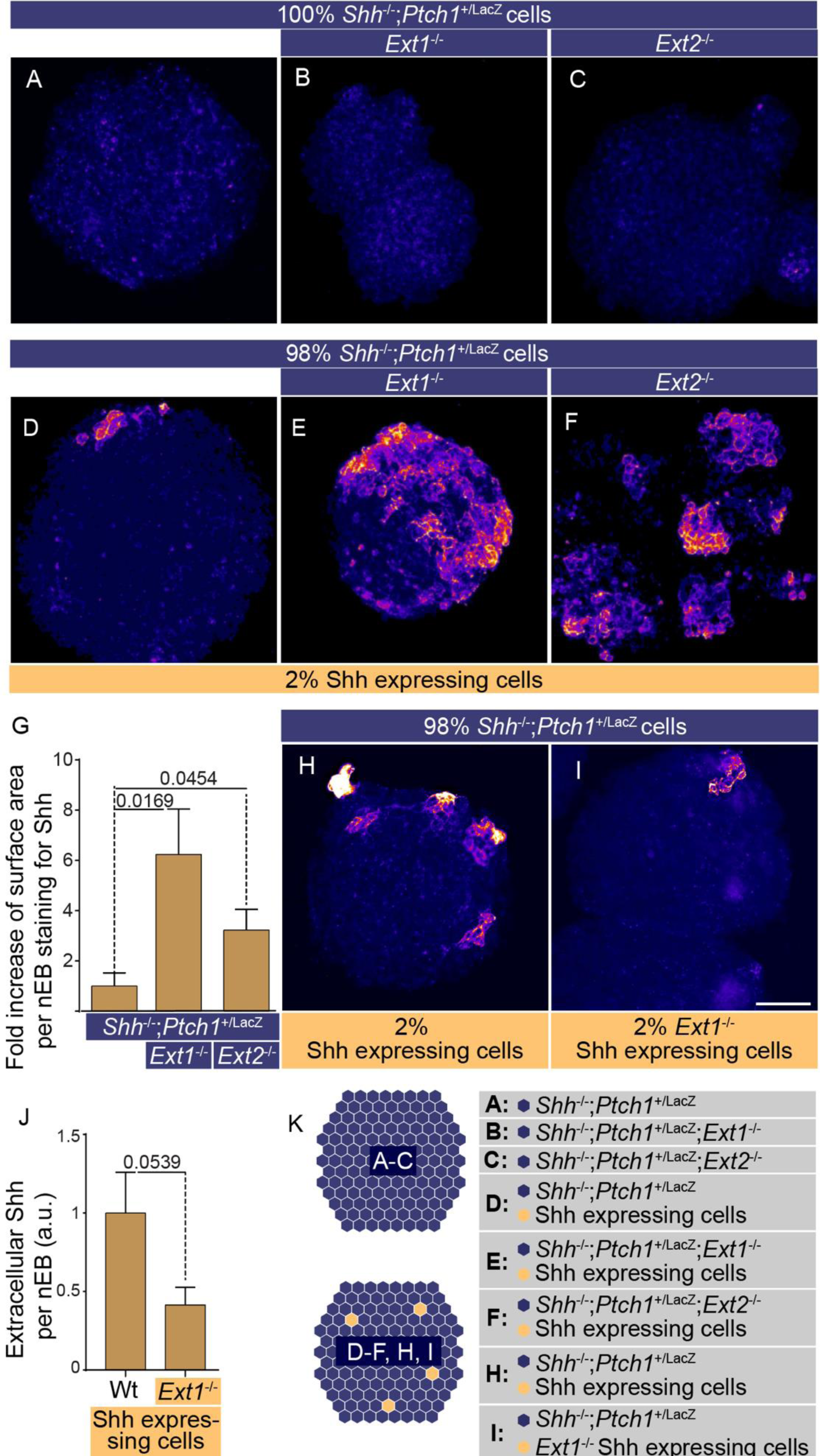
Loss of *Ext1/2* function facilitates distribution of Shh away from its source cells, while loss of *Ext1/2* from the Shh source diminishes Shh accumulation in the extracellular matrix. **A-E, H, I**: Neuralized embryoid bodies (nEBs) derived from mESC lines with the indicated composition and genotype. Extracellular Shh distribution was assayed by live staining after 4 days. nEBs derived from *Shh*^−/−^; *Ptch1*^+/LacZ^ cells (A), *Shh*^−/−^; *Ptch1*^+/LacZ^; *Ext1*^−/−^ cells (B), and *Shh*^−/−^; *Ptch1*^+/LacZ^; *Ext2*^−/−^ (C), cells without embedded Shh-expressing cells. **D-F:** Mosaic nEBs consisting of 98% either *Shh*^−/−^; *Ptch1^+/LacZ^* cells (D), *Shh*^−/−^; *Ptch1*^+/LacZ^; *Ext1*^−/−^-cells (E), or *Shh*^−/−^; *Ptch1*^+/LacZ^; *Ext2*^−/−^ cells (F), incorporating 2% Shh expressing cells, were live stained for Shh. **G:** Quantification of E-F. The area of positive Shh staining per nEB was measured. Mean±s.e.m.; n≥7; *p*-value is indicated; Student’s t-test. **H, I:** Mosaic nEBs consisting either 2% wildtype Shh expressing cells (H), or *Ext1^−/−^* Shh-expressing cells (I) and 98% *Shh*^−/−^; *Ptch1^+/LacZ^* cells were generated and assayed by live staining for extracellular Shh distribution. **J:** Quantification of H and I. The area of positive Shh staining per nEB was graphed, mean±s.e.m.; n≥15; *p*-values are indicated; Student’s t-test. K: Diagram showing the experimental approach. Scale bar is 100µm.

Whereas the loss of *Ext1/2* facilitates distribution of Shh away from the source cells*, Ext1* function appears to enhance Shh presentation of the Shh-expressing cells (Fig. 1 H, I, quantified in J), possibly explaining the negative effects on Shh signaling due to the loss of *Ext1/2*.

### Loss of *Ext1/2* increases the non-cell autonomous response to Shh

As HSPG have been implicated as co-receptors for Shh, we assessed if the loss of *Ext1* affected the response to Shh synthesized at a distal site. We assessed the Shh response by staining for markers of distinct neural progenitor populations along the vertebrate dorsoventral axis in the neuralized nEBs. Olig2 and Isl1/2 served as markers of ventral cell populations that are induced by Shh. Consistent with elevated Shh levels in *Ext1*and *Ext2* null nEBs, but not supporting a role for HSPG as Shh co-receptors, we observe higher levels of Shh-induced Olig2 and Isl1/2 in *Shh*^−/−^; *Ptch1*^+/LacZ^; *Ext1*^−/−^ and *Shh*^−/−^; *Ptch1*^+/LacZ^; *Ext2*^−/−^ nEBs than in *Shh*^−/−^; *Ptch1*^+/LacZ^ nEBs (Fig 2 A-N). As an independent assay for the effects of inactivating Ext1/2 on Shh signaling response, we assessed Ptch1:LacZ induction (Goodrich et al., 1997) in mosaic nEBs. Previous work validated that Ptch1:LacZ induction mirrored ventral neural progenitor differentiation and is a reliable output for Hh pathway activity (Roberts et al., 2016). We observed that Ptch1:LacZ was induced much stronger in *Shh*^−/−^; *Ptch1*^+/LacZ^; *Ext1*^−/−^ and *Shh*^−/−^; *Ptch1*^+/LacZ^; *Ext2*^−/−^ nEBs than in *Shh*^−/−^; *Ptch1*^+/LacZ^ nEBs (Fig. 2O), further demonstrating that inactivation of *Ext1/2* resulted in an enhanced Hh response.

**Figure 2.**
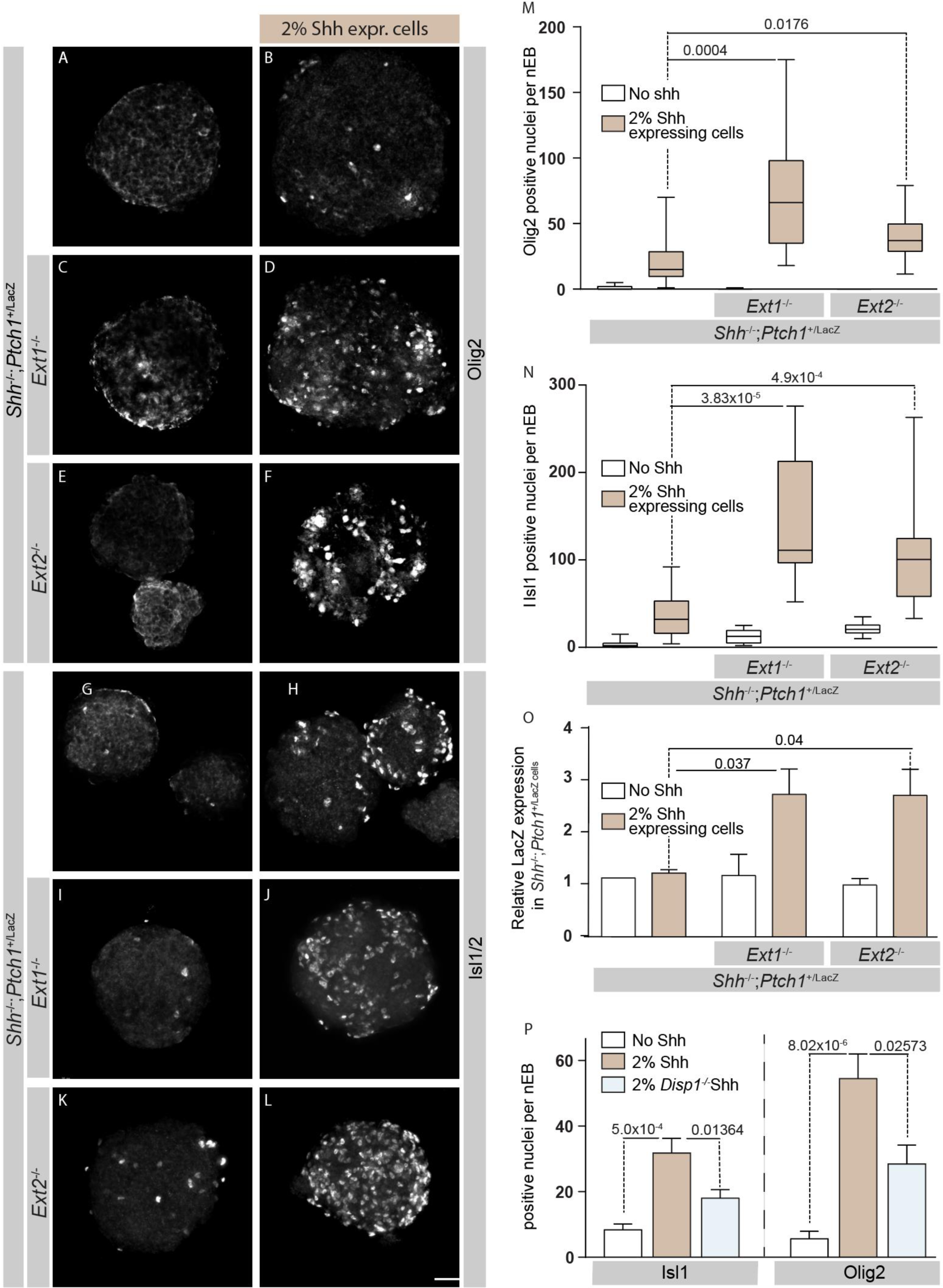
Loss of *Ext1/2* in cells other than the Shh source results in an enhanced Shh response as measured by Isl1/2, Olig2, and *Ptch1* induction. **A-F:** Mosaic nEBs composed of the indicated genotypes and stained for Olig2. **G-L:** Mosaic nEBs composed of the indicated genotypes and stained for Isl1/2. **M:** Quantification of the number of Olig2 positive nuclei per nEB. **N:** Quantification of the number of Isl1/2 positive nuclei per nEB. **O:** The Shh-mediated induction of Ptch1:LacZ was measured in *Shh*^−/−^; *Ptch1*^+/LacZ^, *Shh*^−/−^; *Ptch1*^+/LacZ^; *Ext1*^−/−^, or *Shh*^−/−^; *Ptch1^+/LacZ^*; *Ext2*^−/−^ EBs, and normalized to total protein. Graphed is the mean LacZ activity ±s.e.m.; n=3. **P:** Mosaic nEBs consisting of 2% either wildtype or *Disp1^−/−^* Shh expressing cells and 98% *Shh*^−/−^; *Ptch1^+/LacZ^*; *Ext1*^−/−^ were cultured and assayed for Olig2 and Isl1/2 expression. Data are mean positive nuclei per nEB ±s.e.m.; n≥10; *p*-values are indicated; Student’s t-test. Scale bar is 100µm.

In order to confirm that the enhanced Shh response we observed in *Ext* nulls was due to enhanced Shh distribution from source cells, we blocked Shh release by inactivating *Disp1* in Shh expressing cells. Even though the exact molecular mechanism remains not fully understood, the involvement of *Disp1* in mediating lipid-modified Shh from synthesizing cells is well established (Caspary et al., 2002; Etheridge et al., 2010; Ma et al., 2002). *Disp1* inactivation in Shh expressing cells led to a significant loss of Shh response, revealed by Olig2 and Isl1/2 immunostaining (Fig. 2P), further supporting the notion that the observed changes in response are a direct consequence of Shh distribution. Together, our results indicate that Shh released from cells surrounded by a normal HSPGs preferentially distributes into the extracellular space that lacks the correctly modified HSPGs. Moreover, this increased distribution away from the sites of synthesis results in a significantly enhanced Shh response. This demonstrates that HSPGs negatively affect Shh distribution, and indicates that HSPGs play no obvious role in the presentation of Shh to its cognate receptor.

### *Ext1/2*-dependent HS chains regulate Shh distribution and response non-cell autonomously

Inactivation of *Ext1/2* in nEBs lead to an increased range of Shh distribution and signaling, suggesting that Shh binding to HSPG is inhibitory to Shh function. To address this notion, we treated mosaic nEBs with ectopic heparan sulfate and found that 20 μg/ml HS led to a significant reduction in Olig2 (Fig. 3 A-G) and Isl1/2 (Fig. 3 H-N) induction, indicating that the interaction of Shh with HS restricts its activity.

**Figure 3.**
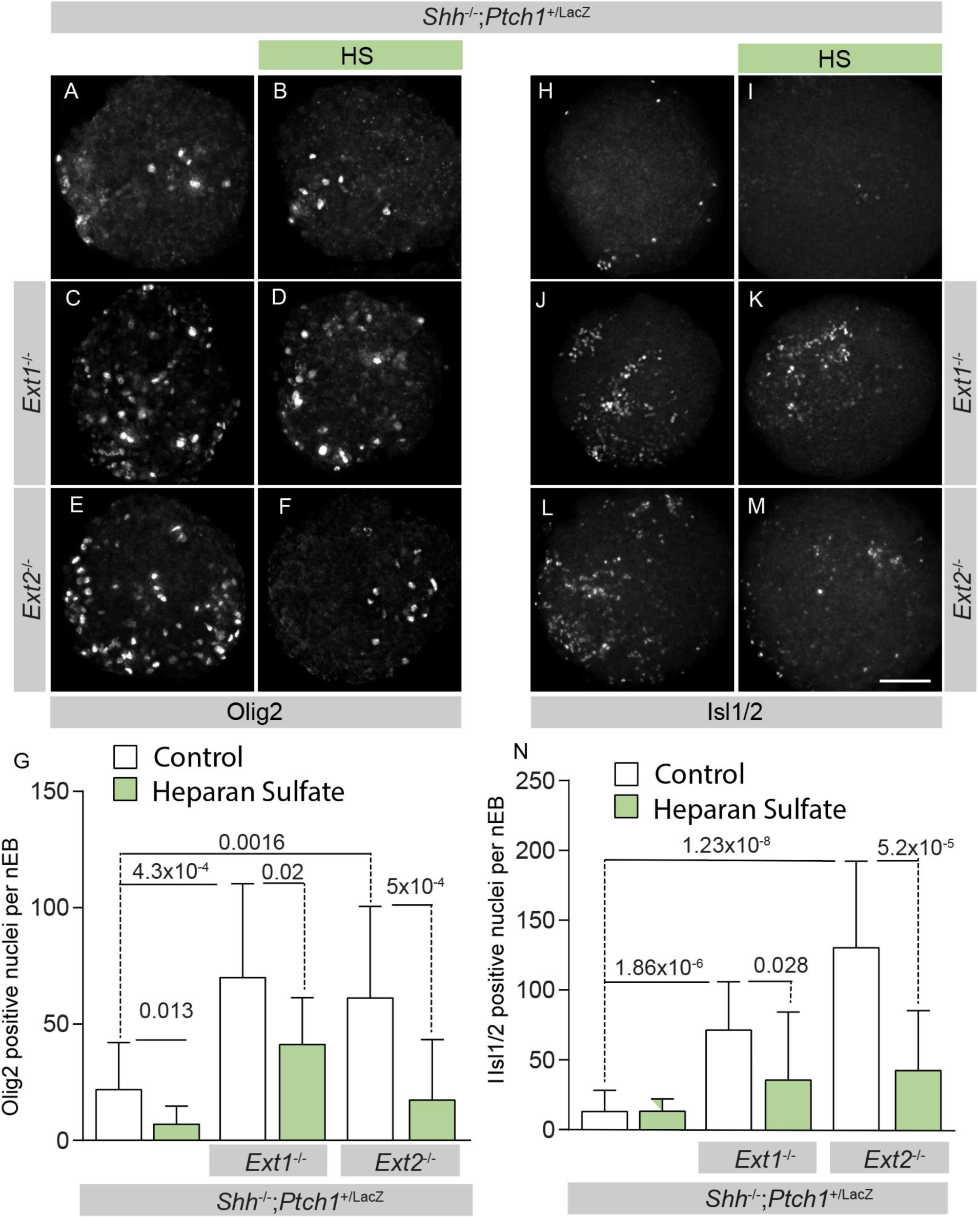
Heparan Sulfate lowers Shh signaling in *Ext1/2* null nEBs. **A-F, H-L**: Mosaic nEBs of the indicated genotypes, including 2% Shh expressing cells. *Shh-*/-; *Ptch1*^+/LacZ^ cells (A, B,H, I) or *Shh*^−/−^; *Ptch1^+/LacZ^*; *Ext1*^−/−^ cells (C, D,J, K) or *Shh*^−/−^; *Ptch1*^+/LacZ^; *Ext2*^−/−^ cells (E, F,L, M) and 2% Shh expressing cells were cultured in control medium (A, C,E, H,J, L) or in medium supplemented with Heparan Sulfate (HS) (B, D,F, I,K, M). **G, N:** Quantification of the Shh-mediated induction of Olig2 (G) and Isl1/2 (N) per nEB, mean±s.e.m.; n≥10; *p*-values are indicated; Student’s t-test.

In order to address the cell-autonomy of Ext1/2-dependent HS chains for Shh signaling, mosaic nEBs comprised of equal numbers of *Shh*^−/−^; *Ptch1*^+/LacZ^ and either *Shh*^−/−^; *Ptch1*^+/LacZ^; *Ext1*^−/−^ or *Shh*^−/−^; *Ptch1*^+/LacZ^; *Ext2*^−/−^ were generated (Fig. 4A). To confirm that HS chains from *Shh*^−/−^; *Ptch1*^+/LacZ^ cells in chimeric nEBs are able to regulate Shh signaling non-cell autonomously, we examined extracellular Shh accumulation in chimeric nEBs. The results indicated that extracellular Shh accumulation was restored to regular level after the introduction of *Shh*^−/−^; *Ptch1*^+/LacZ^ ES cells (Fig. 4B). Moreover, the elevated Shh response in observed in *Ext1/2* null nEBs was suppressed to wild type levels by HSPGs synthesized *Shh*^−/−^; *Ptch1*^+/LacZ^ cells. Mosaic nEBs consisting of equal numbers of *Shh*^−/−^; *Ptch1*^+/LacZ^; *Ext1*^−/−^ and *Shh*^−/−^; *Ptch1*^+/LacZ^; *Ext2*^−/−^ cells have similar high levels of Olig2 and Isl1/2 positive cells as either cell type grown alone (Fig. 4 CD) demonstrating that the mechanism by which the loss of *Ext1* or *Ext2* enhances Shh signaling is similar. The results indicated that restoration within a short-range occurred in HS-deficient cells after the introduction of HS-wildtype ES cells. Therefore, HSPGs play a crucial role in regulating Shh distribution and response in a non-cell autonomous manner.

**Figure 4.**
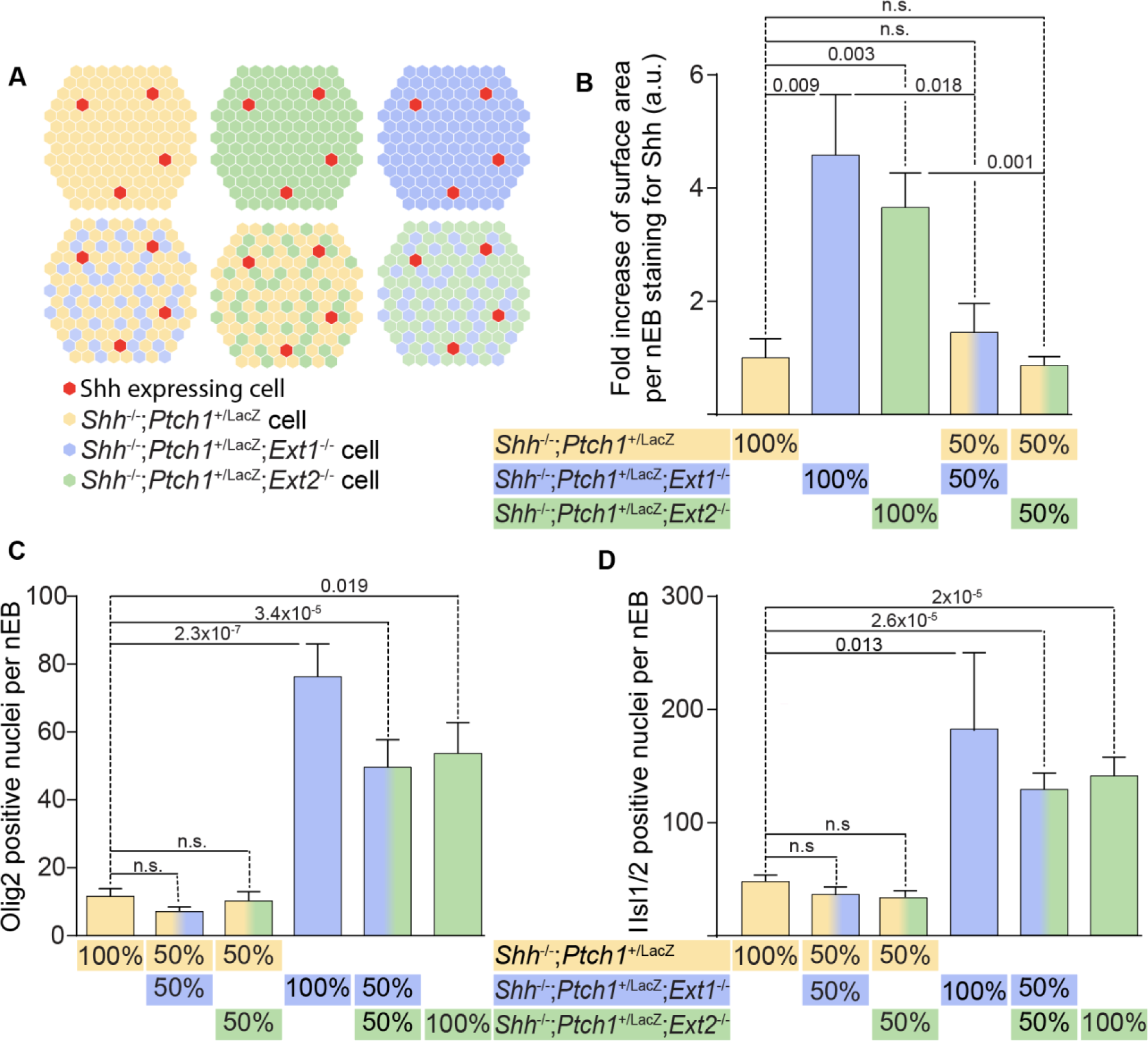
Confined loss of Ext function only in the Shh transporting cells suffices to enhance the Shh response in *Ext1/2* competent cells. **A:** Schematic representations of the experiments. nEBs consisting of 1:1 of *Shh*^−/−^; *Ptch1^+/LacZ cells^*, *Shh*^−/−^; *Ptch1^+/LacZ^*; *Ext1*^−/−^ cells or *Shh*^−/−^; *Ptch1*^+/LacZ^; *Ext2*^−/−^ cells were generated to assess Shh distribution and signaling response. **B:** Quantification of Shh positive area per mosaic nEB composed of the indicated genotypes. Graphed data are mean ±s.e.m.; n=6; *p*-value is indicated; student’s t test. **C, D:** The Shh-mediated induction of Olig2 (C) and Isl1 (D) per nEB was quantified. Data are mean ±s.e.m.; n≥10; *p*-value is indicated; Student’s t-test.

### *Gpc5* is the core protein that is involved in HSPG-mediated Shh distribution and response regulation in nEBs

*Ext1/2* catalyze the glycosylation of multiple distinct HSPG core proteins. To find the specific HSPG that affects Shh distribution we followed an informed approach to identify the required core protein. Three major families of PG core proteins have been characterized: the membrane-spanning syndecans, the glycosylphosphatidylinositol-linked glypicans, and the basement membrane PGs perlecan and agrin (Esko and Selleck, 2002; Yan and Lin, 2009). Glypicans are central for Hh distribution and signaling in *Drosophila* (Filmus and Capurro, 2014). We proceeded to mutate all *Glypican* family members expressed in nEBs, and found that in particular the loss of *Gpc5* led to similar phenotype as *Ext1/2* nulls. We observed that extracellular Shh was increased in *Gpc5* null nEBs (Fig. 5 A-C), and consistent with the loss of *Ext1/2*, we observed that Olig2 and Isl1/2 induction by Shh expressing cells was upregulated in *Gpc5*^−/−^ nEBs as compared in nEBs that are wild type for *Gpc5* (Fig. 5 D-M). To address the sufficiency of Gpc5 to inhibit Shh distribution, a complementation experiment was conducted by creating *Gcp5*^−/−^ mESCs stably expressing Gpc5. The results revealed that *Gpc5* cDNA was able to restore the ability of *Gcp5*^−/−^ mESCs to prevent Shh distribution away from the sites of synthesis and suppress Shh-mediated Olig2 and Isl1/2 induction (Fig. 5 C, N). The similarity in phenotypes between the loss of *Ext1/2* and *Gpc5* indicates that Gpc5 is the core protein that is modified by Ext1/2 to inhibit Shh distribution and signaling response.

**Figure 5.**
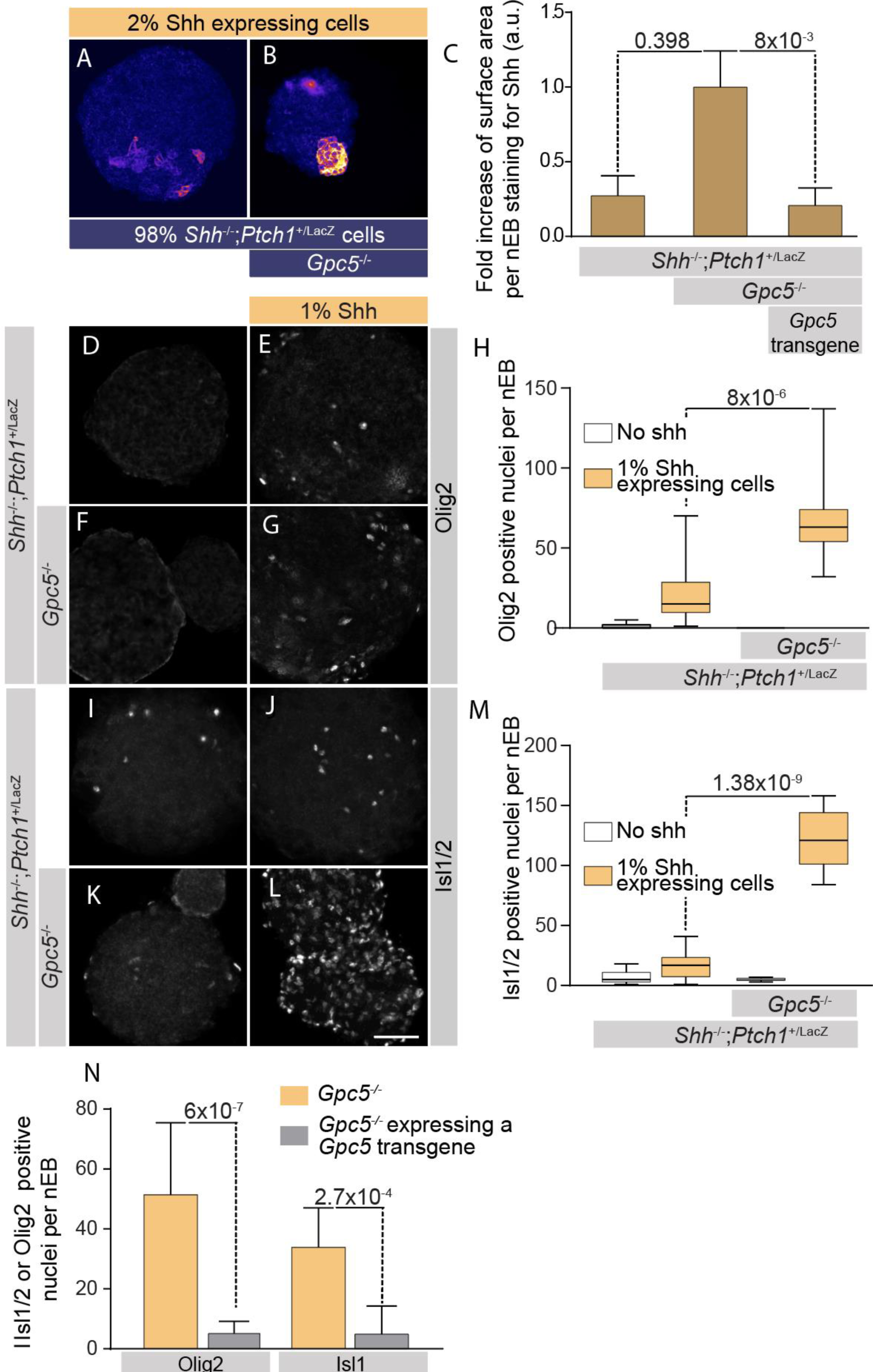
Gpc5 is the core protein that affects HSPG-mediated Shh distribution and response in nEBs. **A, B:** Neuralized embryoid bodies (nEBs) derived from mESC lines with the indicated composition and genotype. Extracellular Shh distribution was assayed by live staining after 4 days. Mosaic nEBs consisting either 98% *Shh*^−/−^; *Ptch1^+/LacZ^* (A) or *Shh*^−/−^; *Ptch1^+/LacZ^*; *Gpc5*^−/−^ (B) and 2% Shh expressing cells were stained for extracellular Shh. **C:** Area of Shh positive per nEB was quantified. Graphed results are the mean±s.e.m.; n≥6, *p*-value is indicated, Student’s t-test. **DL:** Mosaic nEBs consisting either of only *Shh*^−/−^; *Ptch1^+/LacZ^* (D, I) or *Shh*^−/−^; *Ptch1^+/LacZ^*; *Gpc5*^−/−^ (F, K) or incorporated 1% Shh expressing cells (E, G,J, L) were stained for Olig2 (D-G) and Isl1/2 (I-L). **H:** Quantification of D-G. **M:** Quantification of I-K. Graphed is the mean of positive cells per nEB ±s.e.m., n≥10, *p*-value is indicated; Student’s t-test. **N:** A *Gpc5* transgene was stably expressed in *Shh*^−/−^; *Ptch1^+/LacZ^*; *Gpc5*^−/−^ cells. These cells were assessed for their ability to affect the Shh response in mosaic nEBs incorporating 2% Shh expressing cells as compared to the maternal line. The expression of Olig2 and Isl1/2 positive cells per nEB was quantified. Data are mean ±s.e.m.; n≥7, p-value is indicated, Student’s t-test.

### It is sufficient to enhance Shh signaling via disrupting *Ext1/2* or *Gpc5* in the tissue interposed between the Shh source and the responding cells

The molecular mechanism of Shh movement within ECM remains elusive, but the accumulation of Shh that we observed around *Ext1/2* and *Gpc5* null cells indicates that the ECM around cells that lack *Ext1/2* or *Gpc5* is more permeable to Shh. However, it remains a possibility that cells that lack *Ext1/2* or *Gpc5* are intrinsically more sensitive to Shh. To differentiate between these possible explanations, we used a culture system in which we can unambiguously assess the contribution of the cells that transport Shh. We generated tripartite mosaic nEB, consisting of 1) Shh expressing wildtype cells (3% or 5%) as localized Shh sources, 2) 3% of reporter mESCs that have a genetically encoded Shh reporter and are HSPG competent and 3) a predominant compartment of either *Shh*^−/−^; *Ptch1*^+/LacZ^ or cells lacking *Ext1/2* or *Gpc*5, which serve as the conduit for Shh. We used HB9:GFP cells or V3 interneuron reporter Sim1:Cre/tdTomato (Sternfeld et al., 2017) to confine the compartment in which we assess Hh pathway activation. The bulk of the cells in such nEBs is purely assessed for its role in Shh transport between the normal source and responding cells.

Tripartite mosaic nEBs consisting of 97% *Shh*^−/−^; *Ptch1*^+/LacZ^ cells showed negligible Tomato expression (Fig 6 A, B,J, K). In contrast, we observed robust Shh-dependent tomato +V3 induction in mosaic nEBs comprised of 92% *Shh*^−/−^; *Ptch1*^+/LacZ^; *Ext1*^−/−^ or *Shh*^−/−^; *Ptch1*^+/LacZ^; *Ext2*^−/−^ or *Shh*^−/−^; *Ptch1*^+/LacZ^; *Gpc5*^−/−^ cells (Fig. 6 C-I). Similar results were obtained using the HB9:GFP cells as reporters for Shh activity. Tripartite mosaic nEBs consisting of 94% *Shh*^−/−^; *Ptch1*^+/LacZ^; *Ext1*^−/−^ or *Shh*^−/−^; *Ptch1*^+/LacZ^; *Ext2*^−/−^ or *Shh*^−/−^; *Ptch1*^+/LacZ^; *Gpc5*^−/−^ cells had many more HB9:GFP^+^ motor neuron induction than mosaic nEBs principally comprised of*Shh*^−/−^; *Ptch1*^+/LacZ^ cells (Fig. 6 L-R). These results demonstrate that HSPG deficiency in the ECM strongly facilitates Shh distribution between the Shh source and responding cells surrounded by normal ECM, demonstrating that HSPGs negatively affect Shh transport, possibly by Shh sequestration.

**Figure 6.**
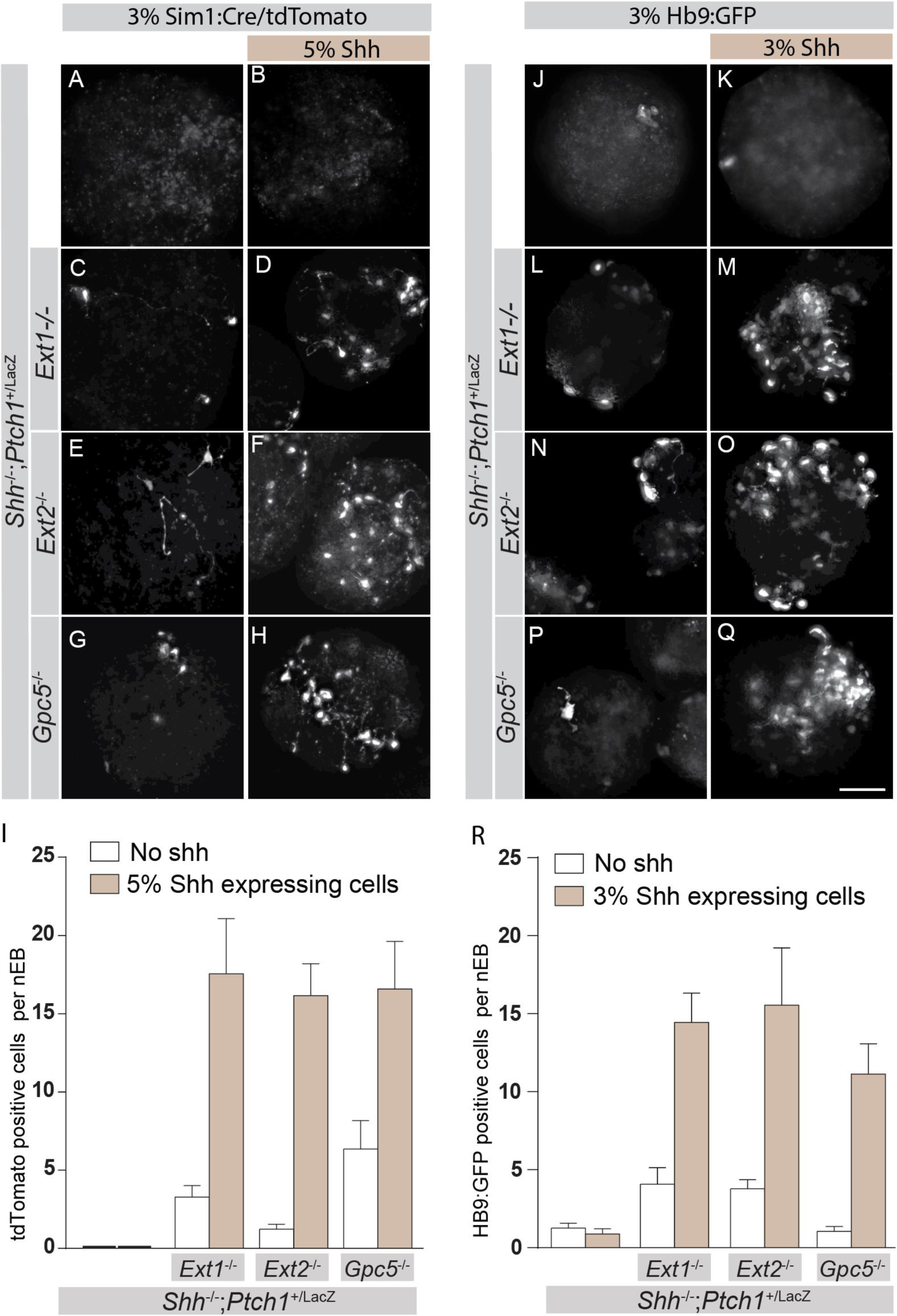
Los of Ext1/2 or Gpc5 only in the Shh transporting cells suffices to enhance the Shh response. **A-H** Neuralized embryoid bodies the bulk of which consists of the indicated genotype as well as 3% of Sim1:RFP cells, with (B, D,F, H) or without (A, C, E, G) 5% Shh expressing cells. Images show RFP expressing cells. **I:** Quantification of A-H. Graphed is the are mean number of positive cells per nEB ±s.e.m., n≥10, *p*-value is indicated; Student’s t-test. **J-Q:** Neuralized embryoid bodies the bulk of which consists of the indicated genotype as well as 3% of Hb9:GFP cells, with (K, M, O, Q) or without (J, L, N, P) 3% Shh expressing cells. Images show GFP expressing cells. **R:** Quantification of J-Q. Graphed is the mean number of positive cells per nEB ±s.e.m., n≥10, *p*-value is indicated, Student’s t-test.

**Figure 7.**
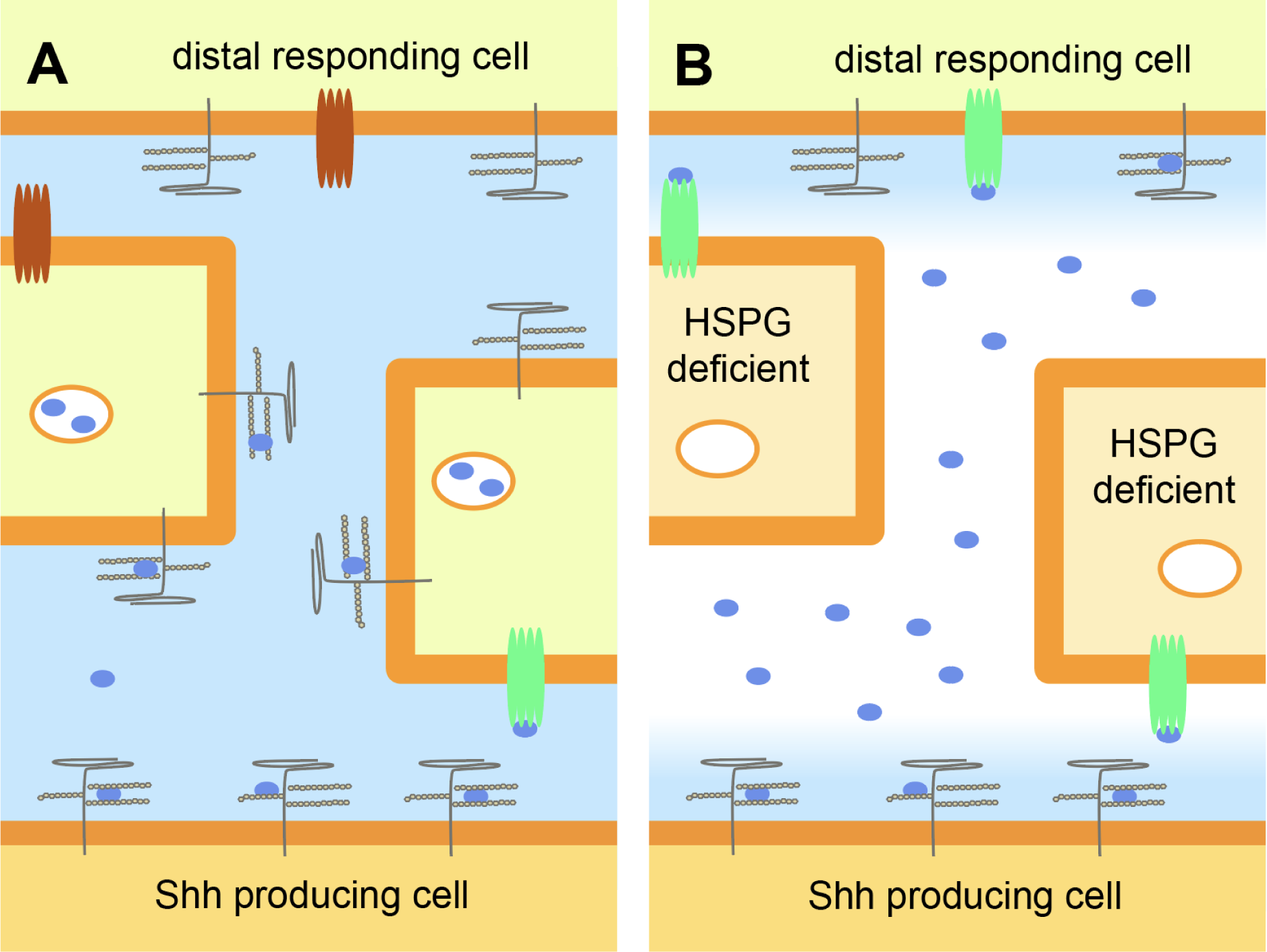
Model. **A:** Under normal conditions the presentation of Shh to the surface of the Shh-expressing cells is facilitated by HSPGs. However, HSPGs prevent widespread distribution of Shh away from the cell presumably by sequestration and endocytosis. **B:** The absence of HSPG form cells other than the Shh expressing cells results in both a wider distribution of Shh and an enhanced Shh response. The presence of HSPG on the responding cells proper does not inhibit the increased Shh response, indicating that HSPGs provide a barrier for Shh thus suppressing paracrine Shh signaling.

## Discussion

In this study, we revealed that *Ext1/2* dependent HS chains regulate extracellular Shh distribution in mosaic nEBs derived from mouse ES cells. Extracellular HS chains play pivotal roles in the local retention of Shh ligands in non-cell autonomous manner, and loss of HS chains in ECM causes dramatic accumulation of Shh and strong upregulation of Shh signaling activity. Our results demonstrate that a central role of Glypicans is to moderate the distribution of Shh, and at least in regard to non-cell autonomous signaling they support earlier notions that Glypicans negatively affect Hh signaling. We find that only in those cells that synthesize and release Shh there is a negative effect on Shh signaling caused by loss of *Ext1*, indicating a facilitating role for HSPGs in Shh signaling.

The roles of HSPG to facilitate Hh signaling was first discovered in *Drosophila*, where loss of either the ext1/2 orthologs tout-velu and sister of tout-velu were found to negatively affect the Hh response. The loss of the Glypican homologs Dally and Dally-like had a similar phenotype compared to the loss of tout-velu. These observations have to be reconciled with the finding that both Ext1/2 and some Glypicans are tumor suppressors, and thus possibly inhibit rather than facilitate signaling.

Shh has a Cardin-Weintraub Motif that mediates its binding to heparan sulfate (Farshi et al., 2011), and HSPGs. This binding could negatively affect Hh signaling by ligand sequestration and limiting distribution (Capurro et al., 2012; Capurro et al., 2008), or it could positively affect Hh signaling by serving as a co-receptor for Hh binding in the target cells (Li et al., 2011). Non-cell autonomous signaling involved three main events: 1) the synthesis and release of the ligand from the source cells, 2) the transport of the ligand through a tissue, and 3) the activation of a receptor in the responding cells. Our results demonstrate that in case of Shh signaling the HSPG function positively affects the presentation of Shh by the source cells, negatively affects the transport of Shh, and has no discernable effect on the ability of cell to respond to Shh, thus providing an explanation for the positive and negative effects of HSPG/Glypicans on Hh signaling. The loss of Shh-sequestration by cells surrounded by an extracellular matrix lacking the proper HSPG complement would provide a simple explanation for why there is more Shh detected in the mutant ECM as well as the enhanced ability to distribute Shh more efficiently. As the overall Hh response increases due to the lack of Ext1/2 or Gpc5 it appears that there is no major role for HSPG in the Shh presentation to its receptor, a notion further supported by our finding that in the tripartite mosaics the normal ECM surrounding the responding cells does not appear to affect the responsiveness.

This enhanced distribution of Shh we observe in tissues surrounded by an ECM that lacks the proper HSPGs would provide an elegant explanation why Ext1 and Gpc5 can function as tumor suppressors. Non-small-cell lung carcinoma often has upregulated Shh expression (Jiang et al., 2015; Vestergaard et al., 2006). As many Shh-induced tumors have distinct Shh-expressing and Shh-transporting/responding compartments it would be advantageous to suppress Glypican expression in the non-Shh-expressing cells. It might be no surprise that in non-small-cell lung carcinomas the loss of Gpc5 function is not uncommon (Guo et al., 2016). Ext1/2 function as tumor suppressors for exostoses, cartilage capped bone tumors that appear next to growth plates. The location is consistent with a role for Ihh that expressed in the growth plate and is required for growth plate maintenance (Robinson et al., 2017) and bone growth. Loss of Ext1/2 would create a domain in which the transport of Ihh is facilitated, and the response unaffected, resulting in the typical bone tumors that characterize somatic loss of Ext1/2 function.

Whereas our findings are generally consistent with the observations of HSPG function in mammals, they are less so with the observations in *Drosophila,* where it appears that Ext1/2 and Glypican facilitate Hh transport away from the source. Although the clonal experiments in the wing disk (The et al., 1999) most closely resemble or experiments using tripartite nEBs, (with a normal Shh source, mutant Shh transporting cells and normal responding cells), we get different results. One explanation for this difference is the reliance in *Drosophila* on cytonemes to distribute Hh (Guerrero and Kornberg, 2014), structures that are not immediately apparent in mammals.

## Materials and methods

### Cell lines

HB9:GFP mESCs were a gift from Dr. Thomas Jessell (Columbia University). Their identity was confirmed by the presence of the Hb9:gfp transgene. Sim1:Cre/tdTomato mESCs were a gift from Dr. Samuel Pfaff (University of California, San Diego). *Shh*^−/−^; *Ptch1*^+/LacZ^ and wild type mESCs overexpressing Shh were previously described (Etheridge et al., 2010; Roberts et al., 2016). mESC lines were maintained using standard conditions without feeder cells.

### Neuralized embryoid body differentiation

mESCs were differentiated into nEBs using established procedures (Wichterle et al., 2002). nEBs were aggregated for 24 hr. in DFNB medium in Petri dishes rotated at 0.8 Hz. 1 µM Retinoic Acid (RA, Sigma, St Louis, MO) was added at 24 hr. nEBs were fixed 72 hr. after the addition of RA for antibody staining of neural progenitors. nEBs were fixed 72 hr. after the addition of RA for imaging and quantifying HB9:GFP fluorescence. Sim1:Cre/tdTomato fluorescence was imaged 96 hr. after the addition of RA.

### Immunostaining

nEBs were fixed with 4% PFA, washed, permeabilized and blocked. The nEBs were then incubated with primary antibodies. For extracellular Shh staining, mouse anti-Shh (5E1) was added to the culture medium 3h before fixation and secondary antibody treatment. Rabbit anti-Isl1/2 was a gift from Dr. Thomas Jessell (Columbia University). Rabbit anti-Olig2 (AB9610) was purchased from MilliporeSigma (St Louis, MO). The samples were then washed and incubated with the appropriate fluorescently labelled secondary antibody (Invitrogen). nEBs were mounted in Fluormount-G and positive nuclei were quantified. Native HB9:GFP and tomato+ fluorescence was imaged directly, after fixation and mounting, without antibody detection. Mounted nEBs were imaged with a Zeiss Observer fluorescence microscope with a 20x objective. Within each experiment, stacks were de-convolved and resulting image files were scrambled for unbiased counting. Images were processed using Fiji ImageJ and Photoshop software (Adobe).

### Genome editing

sgRNAs were designed using the online CRISPR Design tool (http://tools.genomeengineering.org) and cloned into pX459 (Ran et al., 2013). Target sequences of *Ext1* are 5’-TCTTGCCCCACTAAATGGGA-3’ and 5’-GCTTGGGTCCTTCAGATTCC-3’. Target sequences of *Ext2* are 5’-GTTCTATGTAGCAGACAAGC-3’ and 5’-ACTAGATTCCTAGATGGGTA-3’. Target sequences of *Gpc5* are 5’-CGCAAGCCGAACACGAGCCG-3’ and 5’-TGGTGAGACGACGACCTTTC-3’. *Shh*^−/−^; *Ptch1*^+/LacZ^ mESCs were transfected and transiently selected with puromycin. Individual clones were isolated and expanded for further analysis by DNA sequencing of the targeted region. Deletions of sequences were confirmed and sequenced after PCR using primers bracketing the deleted region.

Mention guide seqs here:

### Reporter assays for β-galactosidase

nEBs were collected, washed once in PBS and lysed in lysis buffer [100 mM potassium phosphate (pH 7.8), 0.2% Triton X-100]. Lysates were analyzed using the Galacto-Light chemiluminescent kit (Applied Biosciences, Foster City, CA) for Ptch1:LacZ expression level. Lysates were normalized for total protein using the Bradford reagent (BioRad, Hercules, CA).

## Acknowledgements

This work was supported by NIH grant 1R01GM117090 to HR and a postdoctoral fellowship by Siebel Institute to WG. Rabbit anti-Isl1/2 was a gift from Dr. Thomas Jessell (Columbia University). HB9:GFP mESCs were a gift from Dr. Thomas Jessell (Columbia University). Sim1:Cre/tdTomato mESCs were a gift from Dr. Samuel Pfaff (University of California, San Diego). We also thank Naveen Natesh for his help with generating Ext1/2 sgRNA constructs. WG performed all experiments, WG and HR designed the experiments and wrote the manuscript.

